# *Pseudomonas aeruginosa lasR* Mutants Resist Phagocytosis and Alter Inflammatory Cytokine Production by Cystic Fibrosis Macrophages

**DOI:** 10.1101/2025.09.25.678585

**Authors:** Daniel S. Aridgides, Diane L. Mellinger, Lorraine L. Gwilt, Thomas H. Hampton, Dallas L. Mould, Deborah A. Hogan, Alix Ashare

**Affiliations:** Section of Pulmonary and Critical Care Medicine, Dartmouth-Hitchcock Medical Center, Lebanon, NH; Department of Microbiology and Immunology, Geisel School of Medicine, Hanover, NH; Department of Ecology and Evolutionary Biology, Yale University, New Haven, CT

## Abstract

Cystic Fibrosis is characterized by chronic muco-obstructive lung disease and infection. People with CF (pwCF) are often colonized with *Pseudomonas aeruginosa* for years to decades, allowing for evolutionary adaptation. In chronic *P. aeruginosa* lung isolates from pwCF, the quorum sensing regulator LasR frequently is nonfunctional, however the factors enabling *lasR* loss-of-function (LOF) mutant selection are incompletely understood. We hypothesized that LOF mutations in *lasR* could allow *P. aeruginosa* to resist the selective pressure of phagocytosis. We found that in multiple strain backgrounds, LasR LOF decreased phagocytosis by both model THP-1 and primary monocyte-derived macrophages. While exogenous administration of the quorum-sensing autoinducer 3-oxo-C12-homoserine-lactone (3OC12HSL) that is made by an enzyme regulated by LasR activity inhibited phagocytosis and mitochondrial respiration, the phagocytosis resistance seen with *lasR* mutants appears to be bacterial cell intrinsic rather than due to secreted factors. Finally, we found that *lasR* LOF mutations altered the inflammatory profile upon infection of CF macrophages, with a shift from IL-1 family cytokine expression towards canonical inflammatory markers including IL-6 and TNFα. Collectively these data provide a potential explanation for both the prevalence of *lasR* mutants in the CF lung as well as their association with worse outcomes.

**Importance:** Cystic Fibrosis (CF) is a genetically inherited disease that leads to chronic lung infections. *Pseudomonas aeruginosa* is often implicated with worsening of lung disease, and it evolves in the lung over time to resist eradication. One of the most commonly disrupted gene targets identified in *Pseudomonas aeruginosa* isolates from chronically infected CF lungs is *lasR*, which encodes a transcription factor which regulates multiple virulence factors. What contributes to the apparent fitness of *lasR* mutants in the CF lung is not well known. Our study shows that *lasR* loss-of-function (LOF) mutants resist phagocytosis by macrophages, one of the fundamental mechanisms of clearance by the immune system. We identify mechanisms promoting resistance to phagocytosis, and explore the downstream consequences on inflammatory responses. Understanding why *lasR* mutations arise could inform strategies to eradicate them from the CF lung and improve outcomes.

## Introduction

Cystic Fibrosis (CF) is a multisystem disorder caused by mutations in the Cystic Fibrosis Transmembrane Conductance Regulator (*CFTR*) gene, encoding a chloride and bicarbonate channel (1). In the lungs of people with CF (pwCF), CFTR mutations lead to dehydrated, sticky mucus and subsequent colonization with bacteria often including *Pseudomonas aeruginosa* (2). Novel highly effective modulator compounds which improve trafficking and function of the CFTR protein have markedly improved outcomes for the subset of pwCF harboring CFTR mutations that are responsive to them (3–12), however CFTR modulators cannot restore sterilizing immunity to the lungs, particularly when there is pre-existing structural damage (13, 14). Given their relatively recent availability, the long-term trajectory of lung function in pwCF remains unknown, even for those who are clinically improved on modulators.

Lung function over time is impacted by bacterial colonization, with cycles of infection and dysregulated inflammation leading to lung damage (1). The specific composition of the lung microbiota as well as the host response to them play a role in determining whether lung function worsens or remains stable. Low bacterial diversity, *P. aeruginosa* colonization, and detection of *P. aeruginosa* with disruptions to the LasR/I quorum sensing system have all been shown to correlate with accelerated lung function decline in pwCF (15–17).

LasR loss-of-function (LOF) mutants are one of the most commonly isolated morphotypes found in CF lungs (18). LasR LOF mutants have also been isolated from environments such as pond water, and will arise de novo when *P. aeruginosa* are cultured in vitro (18–21). Their common emergence across various settings argues for a broad (or common) competitive advantage over LasR intact (LasR+) comparators, however the reasons for selection *in vivo* are not entirely clear. Social cheating whereby *lasR* LOF mutants can utilize secreted products from other *P. aeruginosa* isolates without incurring the metabolic cost of producing them has been proposed as a mechanism for *lasR* mutant selection. *lasR* mutation has also been shown increase microoxic fitness and phosphate scavenging (22–24), providing additional possibilities for its evolutionary advantage. Indeed *lasR* LOF mutants are selected for in vitro cultures due to a growth advantage in post exponential phase (25–27) however these experimental systems lack the host component. Interestingly, the phenotype of lack of protease activity has been shown to increase in *P. aeruginosa* isolates that persist in pwCF on highly effective modulators, suggesting *lasR* mutation may also be increased post modulator therapy (28).

In the CF lung *P. aeruginosa* must coexist with host immune cells which are attempting to eradicate it, yet the effects of *lasR* LOF mutations on *P. aeruginosa* interactions with host immune cells remain comparatively unstudied. Studies on interaction with airway epithelial cells (29, 30), neutrophils (31, 32), and macrophages (33) have revealed variable effects on inflammatory responses. Macrophages are responsible for orchestrating the host immune response in the lung, phagocytosis of pathogens and debris, and have been shown to have altered inflammatory and effector activity in CF (34). Although CFTR modulators have been shown to improve CF macrophage function (35–37), an improved mechanistic understanding of how modulators affect the host immune response could benefit the majority of pwCF who are now eligible for highly effective modulator treatment and and lead to novel interventions that might reduce bacterial persistence.

We hypothesized that loss of LasR function would alter interactions with host macrophages as a mechanism of *lasR* mutant persistence within the CF lung, and that CFTR modulators would further impact host response. We found that *lasR* LOF mutants indeed resist phagocytosis in multiple *P. aeruginosa* backgrounds including laboratory and clinical isolates. Phagocytosis resistance was maintained in mixed cultures with LasR+ relatives. *lasR* mutant resistance to phagocytosis by primary CF monocyte derived macrophages (MDMs) was maintained despite the presence of highly effective CFTR modulators. Finally, *lasR* mutation shifted the inflammatory response of MDMs to *P. aeruginosa* infection away from IL-1 family cytokines towards canonical proinflammatory cytokines IL-6 and TNFα. Collectively these results highlight the importance of understanding *P. aeruginosa lasR* LOF in the context of immune cells and provide potential explanations for its increased prevalence and virulence in CF. Some of these results have previously been presented in the form of abstracts (38–42).

## Results

### *lasR* loss-of-function mutations confer phagocytosis resistance independent of background *P. aeruginosa* strain

To test the hypothesis that loss-of-function mutations in *lasR* confer phagocytosis resistance we utilized model THP-1 monocytes which can be differentiated into macrophage-like cells by the addition of the protein kinase C activator phorbol 12-myristate 13-acetate (43). We performed gentamicin-protection assays with three pairs of LasR+ and LasR-*P. aeruginosa* to compare rates of bacterial uptake. We chose a 20 minute timepoint to study phagocytosis as this provides an investigation of uptake alone, whereas longer timepoints may see a mixture of phagocytosis and killing (36). In strain PA14Δ*lasR-*THP-1 co-cultures, fewer cells were phagocytosed then the in the otherwise isogenic wild-type (WT) strain (**Figure 1A**). We then utilized two closely related pairs of LasR+/LasR-clinical isolates for which *lasR* LOF evolved over the course of infection to confirm this effect with *P. aeruginosa* derived from the CF lung. LasR LOF continued to confer phagocytosis resistance in both clinical isolate backgrounds (**Figure 1B,C**) demonstrating that this was not a strain-specific effect.

**Figure 1.**
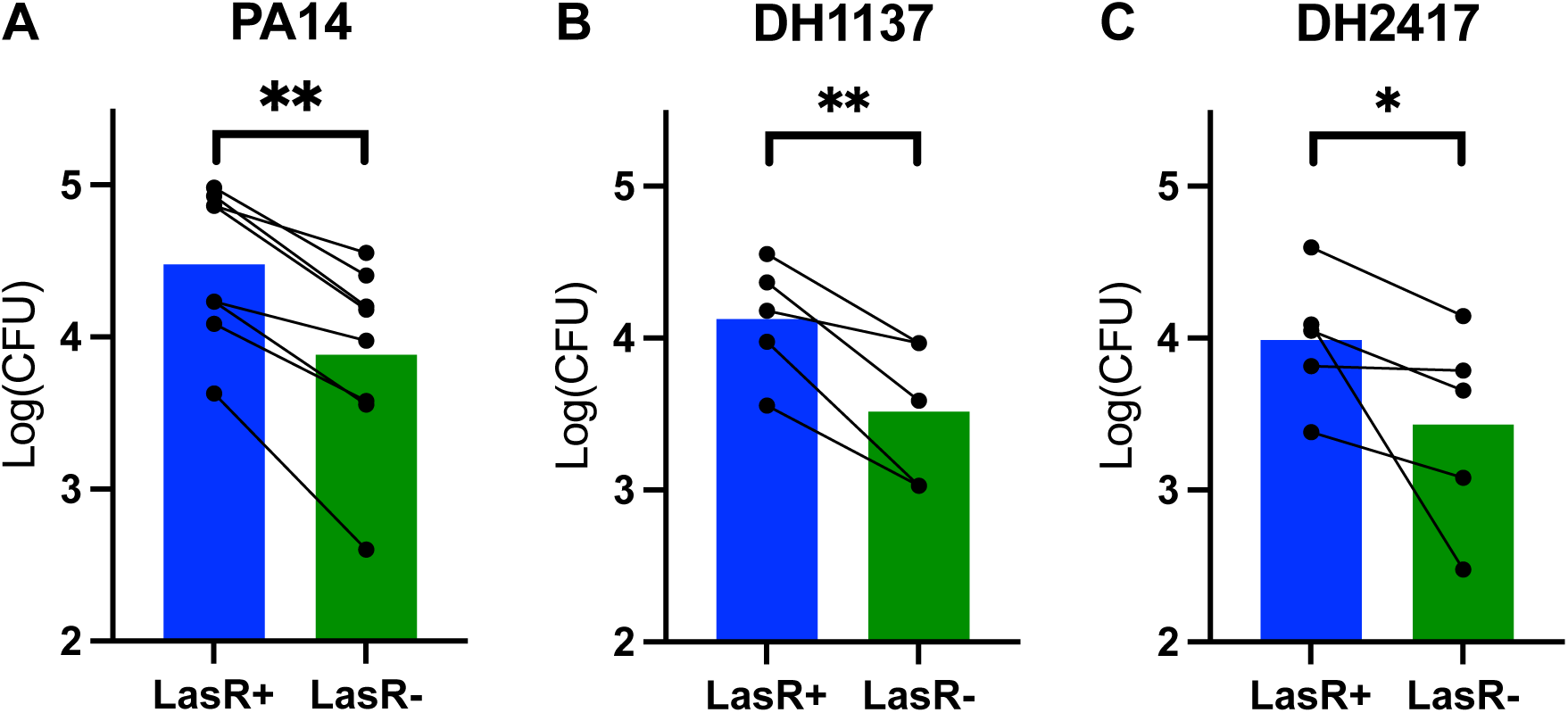
*lasR* mutation confers phagocytosis resistance independent of parental strain. **A)** THP-1 cells were differentiated into macrophages by addition of 50 nM PMA for 48 hours. Phagocytosis was then measured by a gentamicin protection assay at 20 minutes of bacterial uptake. After removal of extracellular bacteria by washing and addition of gentamicin, intracellular CFU were quantified by plating onto LB agar. Phagcoytosis of PA14 wild type and PA14 Δ*lasR* (LasR-) was assessed in eight independent assays, log-transformed means of three technical replicates per assay are displayed. ** p<0.01 for PA14 WT vs LasR-by mixed model linear regression. **B)** Identical assays to **A** were run using naturally occurring paired LasR+/- clinical isolates of *P. aeruginosa* derived from the same parental strain DH1137. ** p<0.01 for PA14WT vs LasR-by mixed model linear regression. **C)** Identical assays to **A** were run using naturally occurring paired LasR+/- clinical isolates of *P. aeruginosa* derived from the same parental strain DH2417. ** p<0.01 for PA14WT vs LasR-by mixed model linear regression.

### LasR but not RhlR autoinducer homoserine lactone inhibits phagocytosis

One of the primary signaling molecules regulated by LasR is the autoinducer N-(3-oxododecanoyl)-L-homoserine lactone (3OC12HSL) that binds to the LasR regulator to induce a positive feedback cycle. 3OC12HSL has been shown to have direct effects on eukaryotic metabolism and signaling (44–55), as well as enhancing phagocytosis of yeast (56). We therefore tested whether 3OC12HSL would increase phagocytosis of *P. aeruginosa* as an explanation for the decreased *lasR* mutant phagocytosis. Contrary to this hypothesis, we found that 3OC12HSL inhibited phagocytosis of both LasR+ and LasR-PA14 at concentrations of 50 μM or higher in a dose-dependent manner (**Figure 2A**). By comparison, the RhlR autoinducer C4-homoserine lactone (C4HSL) had no appreciable effect (**Figure 2B**). Under all concentrations of HSLs tested there continued to be a reproducible difference between LasR+ and LasR-phagocytosis. These results suggest that 3OC12HSL is not responsible for the effects seen in **Figure 1**.

**Figure 2.**
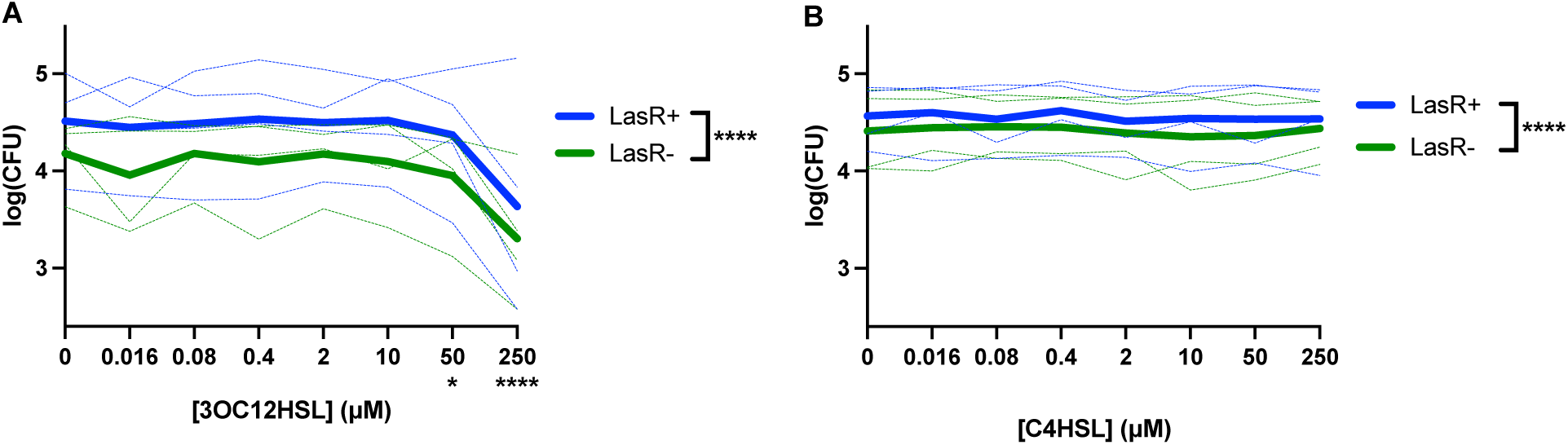
LasR-regulated autoinducer inhibits phagocytosis in a dose-dependent manner. **A)** THP-1 macrophages were incubated with the indicated concentration of 3OC12HSL for 20 minutes prior to performing a gentamicin phagocytosis assay with PA14 WT and PA14Δ*lasR* (LasR-) in parallel. Four separate experiments were performed with three technical replicates each. The means of technical replicates from each individual experiment are shown in the light lines, with darker lines at the means of the four experiments. *p<0.05, **** p<0.0001 for 50 and 250 μM 3OC12HSL relative to DMSO respectively, and LasR+ vs LasR-by mixed model linear regression. **B)** Identical assays to **A** were performed with C4HSL in place of 3OC12HSL. **** p<0.0001 by mixed model linear regression.

### Phagocytosis resistance of LasR-strains is maintained in mixed cultures

We therefore sought to ascertain whether *lasR* mutation effects were due to secreted or cell intrinsic factors. We performed phagocytosis assays with a mixture of LasR+/LasR-PA14 derivatives at varying percentages (0/100, 10/90, 30/70, 50/50, 70/30, 90/10 and 100/0). The LasR-competitor in the PA14 background bore a *lacZ* reporter gene and plating onto X-gal plates allowed for post-assay determination of strain identities when mixed with an untagged wild type. While we found bacterial uptake was generally proportional to the percent input for each strain (**Figure 3A**), there was a persistently lower amount of LasR-*P. aeruginosa* phagocytosed. **Figure 3B** displays the same data as in **3A** graphed relative to percent input for each strain such that the decreased phagocytosis of LasR-strains is evident. These data suggest that the effects of *lasR* mutation on phagocytosis is due to cell intrinsic factors.

**Figure 3.**
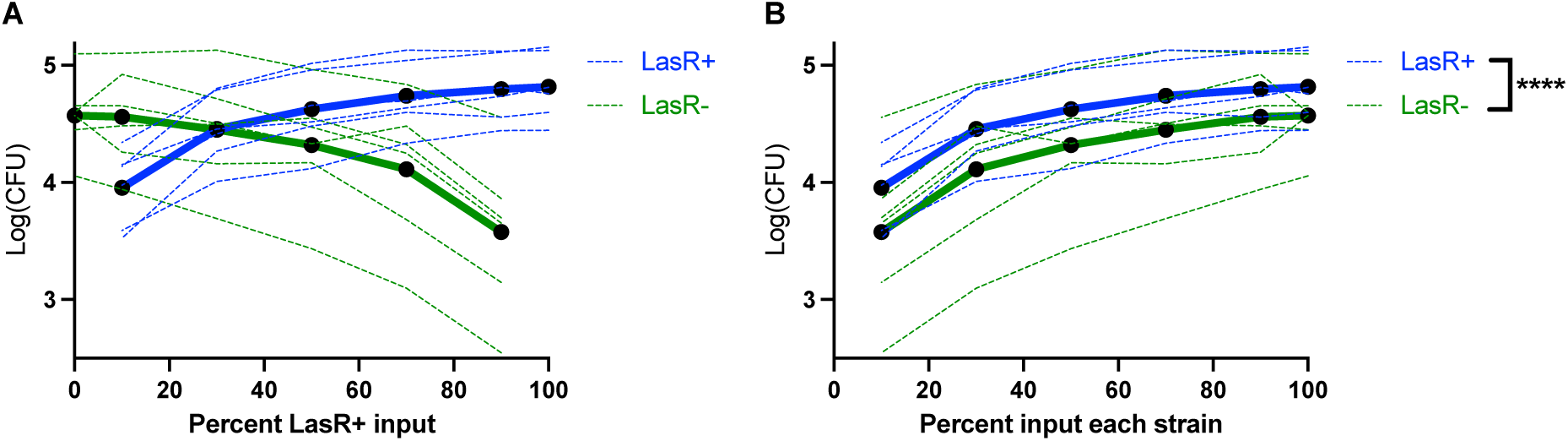
lasR mutation confers phagocytosis resistance in mixed cultures. **A)** A PA14Δ*lasR* strain engineered to express a LacZ reporter was mixed with PA14 wild type at 0%, 10%, 30%, 50%, 70%, 90% and 100% with a fixed total MOI of 10 for the two strains and a phagocytosis assay was performed as previously. After lysis, bacteria were plated onto X-gal plates to allow for differentiation of colony genotype. Means of three technical replicates from 6 independent experiments are graphed, with a dark line at the mean of the 6 experiments. **B)** The same data as in **A** are graphed relative to the percent of each strain input, rather than percent LasR-to better visualize differences. **** p<0.0001 by mixed model linear regression.

### Primary human macrophages augment phagocytosis with CFTR modulators, but LasR phagocytosis resistance is maintained

To determine if the differences between phagocytosis of LasR+ and LasR-cells seen in THP-1 cells were also observed when using primary human cells, we isolated peripheral blood derived mononuclear cells (PBMCs) from volunteers and differentiated them into monocyte-derived macrophages (MDMs) in vitro for seven days. We first tested cells from CF volunteers. We and others have demonstrated that CFTR modulators have a positive effect on macrophage and monocyte phagocytosis (36, 37, 57–63), and as a majority of pwCF are now eligible for CFTR modulators it is important to understand whether they may play an additional role in modifying the response to *lasR* mutants. We tested one pair of laboratory isolates and one pair of clinical isolates (DH1137/6) in MDMs that were treated for 48 hours with CFTR modulators or DMSO as a control. The PA14Δ*lasR* (LasR-) strain was less efficiently phagocytosed by primary CF MDMs relative to its LasR+ progenitor (**Figure 4A**). Tezacaftor/ivacaftor (TI) and elexacaftor/tezacaftor/ivacaftor (ETI) improved phagocytosis of both strains in the PA14 background (**Figure 4A**) however there was no statistically significant interaction with *lasR* status. We found similar results using paired clinical *P. aeruginosa* isolates, where *lasR* mutants were less efficiently phagocytosed and CFTR modulators improved phagocytosis of both strains with no difference in response to modulators between isolates (**Figure 4B**). We have previously demonstrated that nonCF MDMs respond similarly to CFTR modulators when compared with CF MDMs (37), therefore we tested whether any of the effects on *lasR* mutant phagocytosis might be modified by *CFTR* genotype. NonCF MDMs displayed a similar pattern, with decreased phagocytosis of *lasR* mutants, improved phagocytosis in the presence of CFTR modulators, and no statistically significant interaction between modulators and *lasR* mutation (**Figure 4C**). With the LasR+/LasR-clinical isolates, there continued to be an effect of *lasR* mutation (**Figure 4D**) however double CFTR modulator treatment only displayed a non-significant trend towards enhanced phagocytosis, with triple modulators still proving effective (**Figure 4D**). Collectively these results confirm that *lasR* mutants exhibit reduced phagocytosis by primary human CF and nonCF macrophages, and that while CFTR modulator treatment is still beneficial, it fails to restore phagocytosis to the level of LasR+ strains.

**Figure 4.**
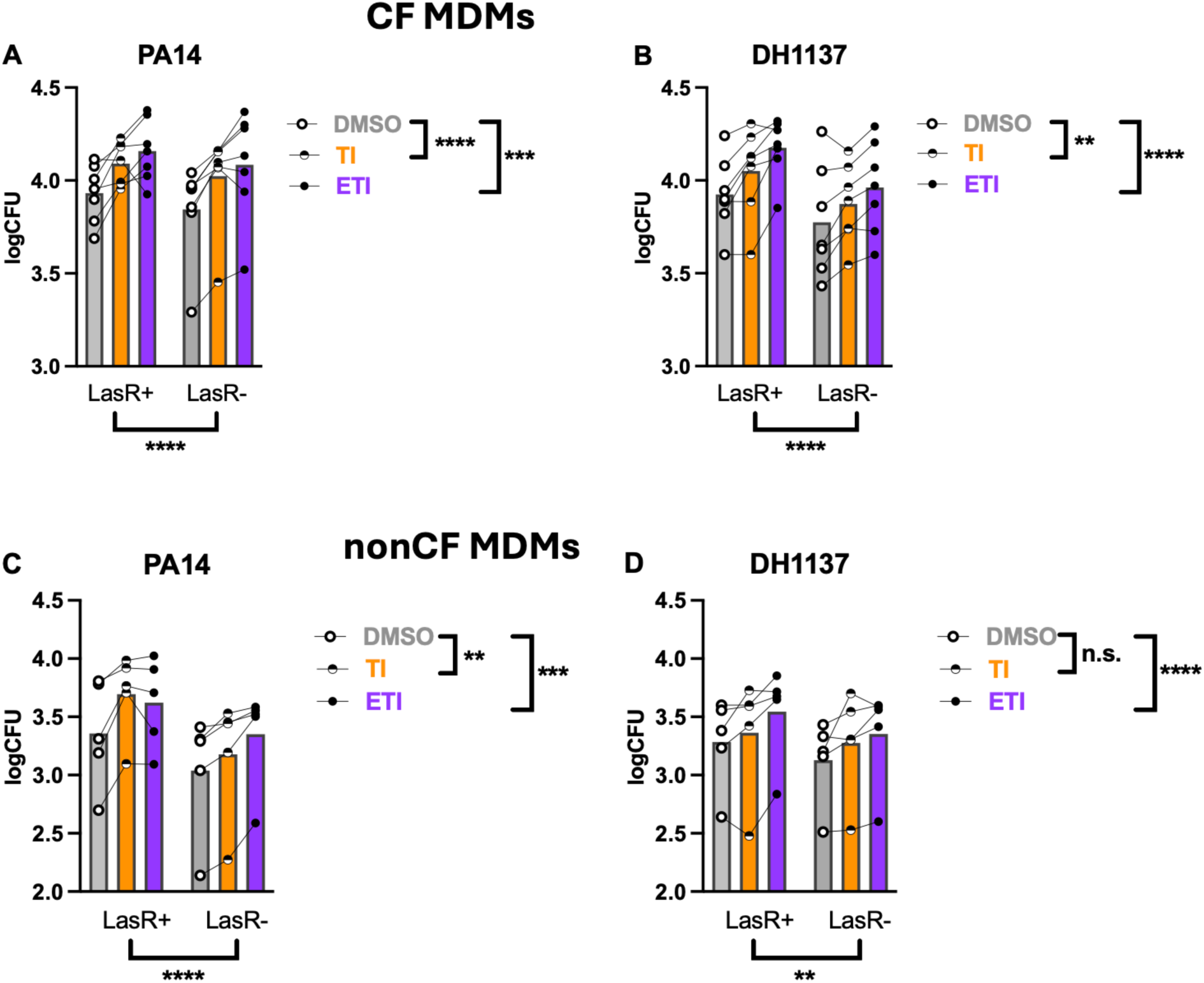
*lasR* mutants exhibit reduced phagocytosis by primary MDMs irrespective of CFTR genotype or modulators. **A)** CF MDMs were treated for 48 hours with CFTR modulators as indicated, then infected with PA14 WT or PA14 *ΔlasR* (LasR-). Lines connecting dots represent data from a single subject (n=7). **B)** As in **A** except the clinical isolate DH1137 and its derivative *lasR* mutant were used. **C)** NonCF MDMs (n=5) were infected with PA14 WT or LasR-derivative as in **A**. D) NonCF MDMs were infected with DH1137 or its *lasR* mutant derivative as in **B**. ** p<0.01, ***p<0.001, **** p<0.0001 for indicated comparisons by mixed model linear regression. Data were combined with previously published data on LasR+ strains only (37).

### *lasR* mutation abrogates the inhibitory effect of *P. aeruginosa* conditioned medium on macrophage mitochondrial respiration

Macrophage metabolism impacts their inflammatory predilection, with activated macrophages relying on glycolysis for ATP generation whereas pro-resolution utilized aerobic respiration (64, 65). We have previously demonstrated that secreted products from *P. aeruginosa* inhibit aerobic respiration in MDMs (37), and we hypothesized that *lasR* mutation may impact this effect. In addition, we saw that ETI inhibits aerobic respiration without significantly affecting glycolysis. We therefore performed Seahorse metabolic flux analysis on CF MDMs pre-treated with CFTR modulators and then then conditioned media from either LasR+ or LasR-PA14 strains was added acutely. Oxygen consumption rate (OCR) was measured as a proxy for mitochondrial respiration and extracellular acidification rate (ECAR) as a proxy for lactate generation/glycolysis. **Figure 5A-C** represent example readouts of OCR from a single CF subject pre-treated with DMSO, TI or ETI, respectively and **Figure 5D-F** are the corresponding readouts for ECAR. Addition of oligomycin, FCCP and rotenone/antimycin A allow for measurement of additional parameters related to mitochondrial respiration. In CF MDMs, ETI decreased basal respiration similarly to prior experiments whereas TI had no effect (**Figure 5G**). Upon acute injection of LasR+ conditioned medium, there was a decrease in mitochondrial respiration (**Figure 5H**), however LasR-conditioned medium completely lost this effect. In the setting of low basal mitochondrial respiration with ETI, the effect of *P. aeruginosa* conditioned medium (either LasR+ or LasR-) was lost (**Figure 5H**). Corresponding decreases in ATP-linked respiration were seen with LasR+ conditioned medium and ETI pre-treatment (**Figure 5G**). Basal glycolysis was not affected by CFTR modulators or different prior to injection (**Figure 5J**) however LasR+ but not LasR-conditioned medium caused an acute rise in glycolysis (**Figure 5K**). This effect was lost upon ETI pre-treatment (**Figure 5K**). OCR/ECAR ratio is a way of gauging relative contributions of each pathway to ATP-generation, and in line with the effects seen on ATP-linked respiration and glycolysis, OCR/ECAR was decreased with LasR+ but not LasR-conditioned media, as well as by ETI (**Figure 5L**). Measurements of maximal respiration, spare respiratory capacity, non-mitochondrial respiration and proton leak are shown in **Supplemental Figure 1**. These results collectively demonstrate that *lasR* mutation eliminates the mitochondrial effects of *P. aeruginosa* exoproducts. Additionally, CFTR modulators inhibit basal mitochondrial activity and there is no additive effect of *P. aeruginosa* exoproducts. Both of these observations have potential downstream implications for induction of inflammation.

**Figure 5.**
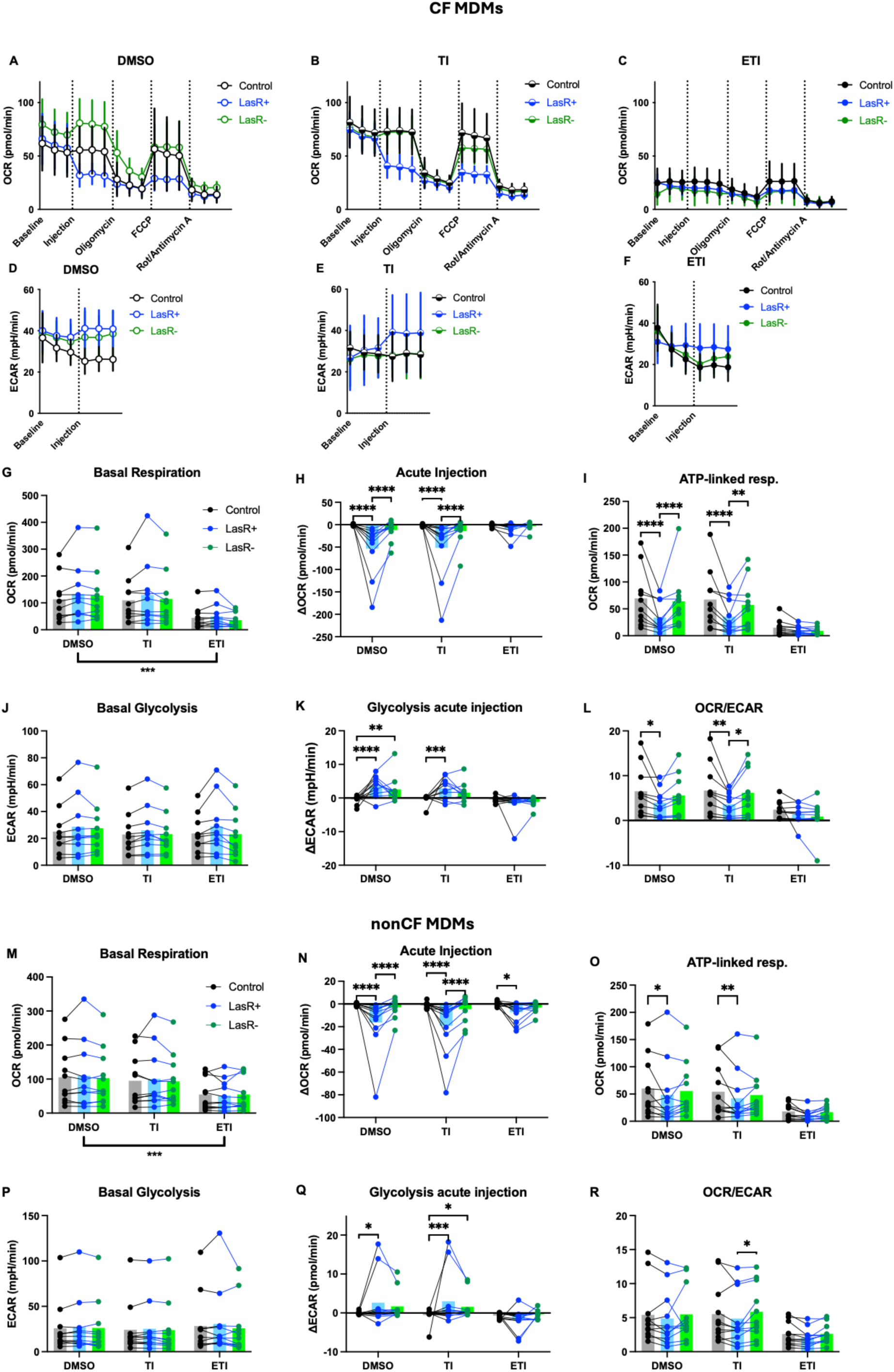
Effects of CFTR modulators and LasR+/- secreted products on macrophage metabolism. CF and nonCF MDM were differentiated with M-CSF then treated for 48 hours with CFTR modulators. A Seahorse mitostress assay with acute injection of *P. aeruginosa* conditioned medium was then performed. **A-C**) Representative oxygen consumption rate tracings from a CF subject MDMs pretreated with DMSO, TI, or ETI. **D-F**) Representative extracellular acidification rates from the same subject. **G-L)** Calculated parameters as indicated based on the mitostress assay of CF MDMs (n=10). Each point with connecting lines represents the mean of technical replicates from an individual subject. Bars represent means of all the subjects within the group. **M-R**) Calculated parameters as indicated for nonCF MDMs (n=11). *p<0.05, ** p<0.01, ***p<0.001, **** p<0.0001 for indicated comparisons by mixed model linear regression. Data were combined with previously published data on LasR+ strains only (37).

We next examined nonCF MDMs to determine whether there may be any differences in how they respond to *lasR* mutants. Similarly to CF MDMs, we found the ETI inhibits basal respiration (**Figure 5M**) with no effect of TI, in agreement with our prior published work (37). LasR+ but not LasR-conditioned medium inhibited mitochondrial respiration (**Figure 5N**) and decreased ATP-linked respiration (**Figure 5O**). These effects were decreased or eliminated by ETI-pretreatment. Modulators had no effect on basal glycolysis (**Figure 5P**), with increased glycolysis secondary to LasR+ but not LasR-conditioned medium (**Figure 5Q**). In contrast to CF MDMs, differences in OCR/ECAR were not statistically significant (**Figure 5R**). Measurements of maximal respiration, spare respiratory capacity, non-mitochondrial respiration and proton leak are shown in **Supplemental Figure 2**.

### LasR autoinducer homoserine lactone inhibits macrophage mitochondrial respiration and induces glycolysis

The divergence between LasR+ and LasR-conditioned medium effects on MDM metabolism led us to hypothesize that 3OC12HSL might underlie these effects. To test this, we returned to THP-1 derived model macrophages and performed Seahorse metabolic analysis with acute injection of 3OC12HSL, with C4HSL as a control. After first confirming no differences in basal respiration prior to injection (**Figure 6A**), we found a dose responsive effect of 3OC12HSL injection (**Figure 6B**) causing inhibition of mitochondrial respiration whereas C4-HSL had no effect. This was partially reflected in differences in ATP-linked respiration (**Figure 6C**) although the effects were not statistically significant. Simultaneous measurement of glycolytic rates confirmed no basal differences in glycolysis (**Figure 6D**), with a dose-dependent increase in glycolysis upon 3OC12HSL injection (**Figure 6E**) and no effect of C4HSL. The shift in metabolic profile from mitochondrial respiration to glycolysis by 3OC12HSL was confirmed with decreased OCR/ECAR (**Figure 6F**). Measurements of maximal respiration, spare respiratory capacity, non-mitochondrial respiration and proton leak are shown in **Supplemental Figure 3**. These results suggest that 3OC12HSL may underlie the effects of LasR+ *P. aeruginosa* exoproducts on mitochondrial respiration.

**Figure 6.**
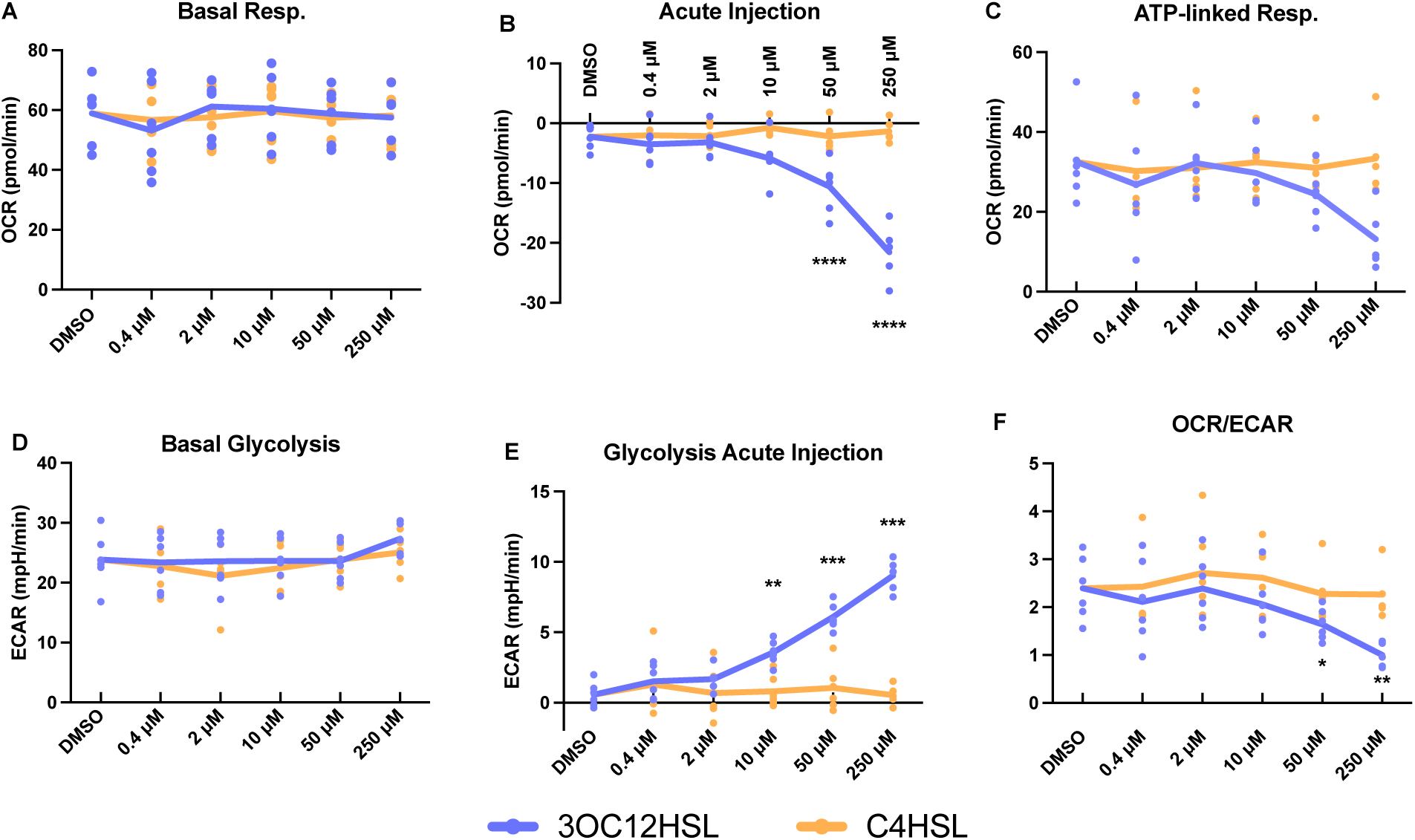
3OC12HSL inhibits mitochondrial respiration in THP-1 macrophages. **A-F)** Seahorse mitostress assay was run with acute injection of various concentrations of 3OC12HSL or C4HSL. Parameters were calculated as per standard protocol. Points represent averages of 6-8 technical replicates from 6 individual experiments, with solid lines at the global means. *p<0.05, ** p<0.01, ***p<0.001, **** p<0.0001 for indicated comparisons relative to DMSO treated wells by mixed model linear regression.

### *lasR* mutation shifts inflammatory cytokine profiles of primary human macrophages away from IL-1 family cytokines towards IL-6 and TNFα irrespective of CFTR modulators and *CFTR* mutation status

Finally, we investigated whether *lasR* mutation impacted inflammatory profiles of MDMs, and the interplay with CFTR modulators. We pretreated MDMs with CFTR modulators for 48 hours, then infected MDMs for two hours with LasR+ or LasR-clinical isolates of *P. aeruginosa* and performed cytokine multiplex analysis on the supernatants. Unsurprisingly we found broad induction of multiple inflammatory cytokines upon *P. aeruginosa* infection of CF MDMs (**Figure 7A**). There was little appreciable effect of CFTR modulators, consistent with our prior work and others’ (37, 66). NonCF MDMs had a similar inflammatory response to CF MDMs, with no statistically significant differences between the groups (**Figure 7B**). We then combined results and tested specifically for differences between LasR+ and LasR-infection. IL-1 family cytokines IL-1α, IL-1β and IL-18 were higher with LasR+ infection, whereas IL-6, IL-10, MCP-1, MIP-1α, and MIP-1β were higher with LasR-infection (**Figure 7C**), demonstrating a specific alteration in the inflammatory profile after macrophage response to *lasR* mutants.

**Figure 7.**
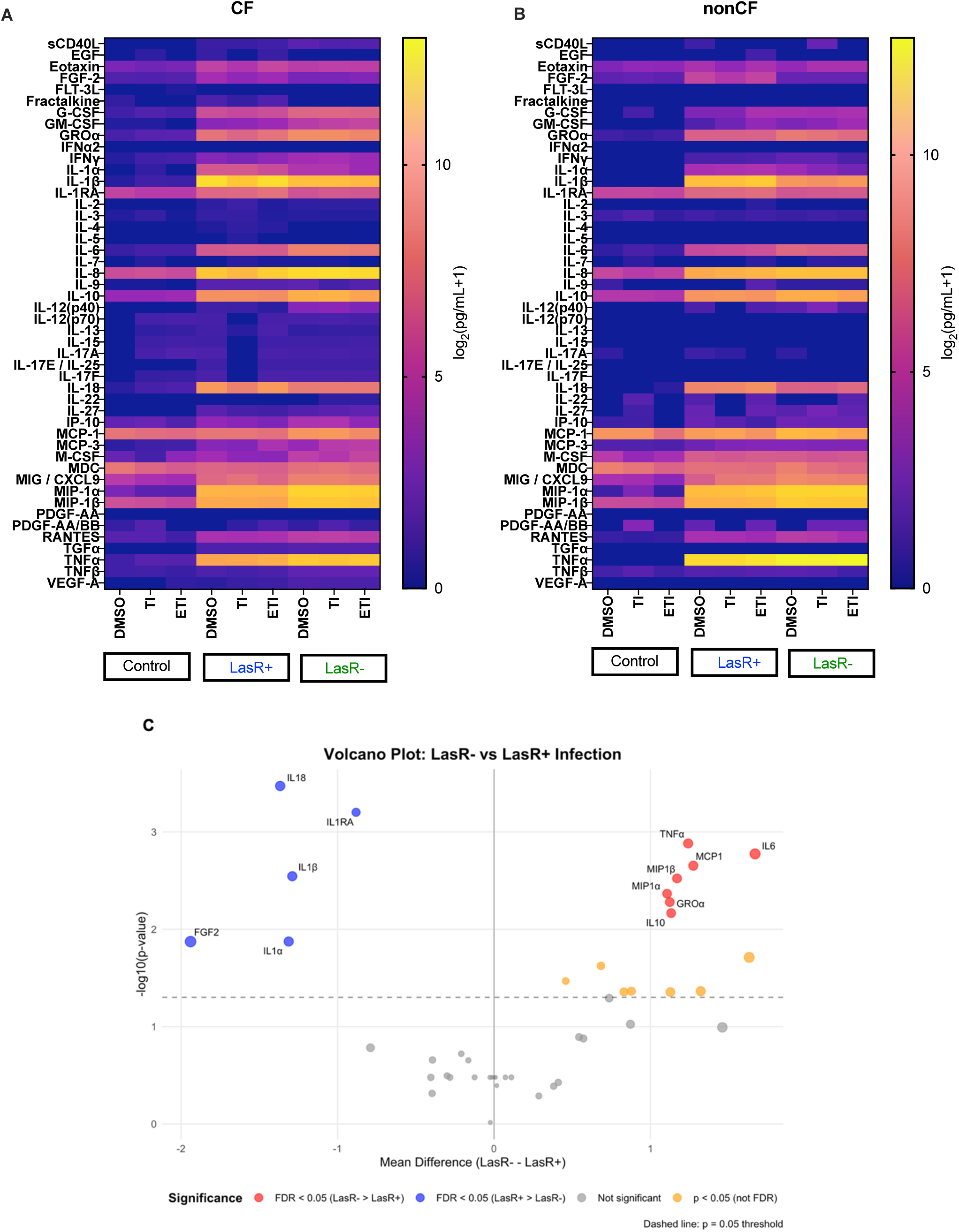
*lasR* mutation alters inflammatory cytokine expression profile by CF and nonCF MDMs. **A)** CF MDMs (n=11 subjects) were pre-treated with CFTR modulators and then infected with DH1137 (LasR+) or DH1136 (LasR-) for two hours, then cells were washed and media replaced. Supernatants were collected 2 hours after washing and analyzed by cytokine multiplex assay. Means of log-transformed data are graphed. **B)** nonCF MDMs (n=4) were infected in the same way and cytokines analyzed by multiplex. C) Volcano plot of fold difference between LasR+ and LasR-cytokine results, DMSO treated group only.

## Discussion

Loss-of-function mutations in *lasR* have been demonstrated to be important for *P. aeruginosa* persistence and virulence within the CF lung (15, 18). How this impacts interactions with host immune cells and specifically in the presence of highly effective CFTR modulators is incompletely understood (29–33, 67). We have identified for the first time a robust and reproducible resistance to phagocytosis by macrophages that is hypothesized to contribute to persistence in the CF lung. While the difference was modest (0.3 to 0.5 log depending on the assay), this represents a relative difference of 2-3 fold uptake in 20 minutes, therefore over years this could represent a significant evolutionary pressure. The phagocytosis resistance was present in multiple background strains including a common laboratory strain (PA14) and two pairs of clinical isolates from the lungs of pwCF (**Figure 1**), increasing the generalizability of our findings.

We also identified potential factors by which LasR mutants resist phagocytosis. The LasR autoinducer 3OC12HSL inhibited *P. aeruginosa* phagocytosis (**Figure 2**), in contrast to prior studies of yeast phagocytosis (56). If 3OC12HSL were responsible for differences in phagocytosis however, it would be expected that LasR mutants would exhibit *higher* phagocytosis rates, which was the opposite of what was seen. Differences in phagocytosis rates were also maintained between LasR+ and LasR-strains when they were mixed in different ratios (**Figure 3**), arguing that the responsible factors are cell intrinsic rather than secreted.

We also explored the interplay between CFTR modulators and *lasR* mutation with phagocytosis and inflammation in primary MDMs. While CFTR modulators were able to enhance phagocytosis of all strains tested consistent with our prior work (37), the decrement with *lasR* mutation was maintained (**Figure 4**). Given the known persistence of *P. aeruginosa* in the lungs of pwCF despite highly effective modulator therapy (14), this is relevant to understanding *P. aeruginosa* colonization. We previously saw that *P. aeruginosa* secreted products inhibited mitochondrial respiration in MDMs (37), as did the triple modulator combination ETI. Intriguingly we found that *lasR* mutation essentially eliminated this effect (**Figure 5**). In combination with the effects of exogenous 3OC12HSL (**Figure 6**), this suggests that the acyl-homoserine lactone is the responsible factor, in line with previous studies of epithelial cells and fibroblasts (45, 46, 48–51, 68).

Unsurprisingly given the variable rates of phagocytosis and differences in effects on mitochondrial function in MDMs, we found that *lasR* mutation impacted inflammatory cytokine profiles upon infection of CF and nonCF MDMs (**Figure 7**). While infection generally induced a broad inflammatory cytokine profile, there was a shift from IL-1 family cytokines IL-1α, IL-1β, IL-1RA and IL-18 towards more classical inflammatory markers IL-6 and TNFα, as well as macrophage inflammatory protein 1α, 1β and macrophage chemoattractant protein 1. There was a trend towards increased IL-8 for *lasR* mutant infection that was not statistically significant after adjustment for multiple comparisons (**Figure 7C**). A shift towards IL-1 family cytokines may reflect increased inflammasome activation versus direct proteolytic activation of IL-1 family precursor proteins. The LasR-regulated elastase LasB has previously been demonstrated to directly activate pro-IL-1β in mice and human macrophages (69, 70), consistent with our data showing less mature IL-1β for infections with *lasR* loss-of-function mutants (**Figure 7C**). Early CF lung *P. aeruginosa* isolates have been shown to be more potent inflammasome activators when compared to later isolates (71), however LasR functional status was not tested in the study by Phuong et al. Decreased protease expression has been seen in persistent *P. aeruginosa* isolates in those on ETI, also suggesting *lasR* mutation may be increased post modulators (28). It is important to note that in our system significant LasR quorum sensing activation would not necessarily be expected, as bacteria are sub-cultured at low density prior to addition to cells, and cells were infected for 2 hours. In contrast to its effects increasing processed IL-1β, LasΒ can decrease IL-8 levels upon infection of airway epithelial cells via direct digestion of IL-8 (72). Future experiments will help to elucidate the exact mechanism whereby *lasR* mutation alters the inflammatory cytokine milieu in the shorter term. We are also currently conducting transcriptomic studies on macrophages infected in parallel with LasR+ or LasR-isolates which should shed light on whether the effects are transcriptional versus post-transcriptional.

Strengths of our work include analysis of *lasR* mutations in multiple backgrounds including a common laboratory strain as well as naturally occurring clinical isolates to improve the generalizability. We began to investigate the mechanisms by which *lasR* loss-of-function mutants may confer phagocytosis resistance although work remains to further elucidate the full mechanism. We confirmed the results found in a human cell line (THP-1 cells) with primary human macrophages from multiple subjects and investigated the interplay with CFTR modulators and genotype (mutant vs wild-type CFTR). Studies performed in parallel on cells from the same subject allowed for a close interrogation of the effects of CFTR modulators and LasR mutation on phagocytosis, metabolism and inflammation.

Our work also carries several limitations. MDMs, while they are a commonly used surrogate for primary lung macrophages due to their ease of procurement, do not behave identically to lung macrophages (73). We plan to expand our work with primary lung macrophages isolated via bronchoscopy (74), and compare them with classically generated MDMs as well as the novel differentiation protocol proposed by Pahari et al. Additionally, while our ex vivo studies provided the opportunity to directly compare effects of LasR status and modulators in paired samples from the same subject, they are necessarily reductive and cannot replicate in vivo conditions. Mouse models differ from human CF in chronicity and acuity of lung infection, however there are data suggesting *lasR* mutants are more virulent in mice as well (31, 72). Finally, we found no direct effect of CFTR modulators including ETI on inflammatory responses by MDMs, in contrast to human studies pre- and post-modulator which demonstrate decreased inflammation (14, 60, 60, 75–78). In vivo studies however cannot separate direct effects on immune cells from decreased mucus and pathogen burden leading to a systemic decrease in inflammation.

In summary, *lasR* mutation confers reproducible resistance to phagocytosis, altered metabolic effects on macrophages, and a change in inflammatory cytokine repertoire upon infection. These effects were independent of background strain of *P. aeruginosa*, as well as CFTR genotype and CFTR modulator therapy. Our work adds to the body of literature on how and why *lasR* mutants are problematic in the lungs of pwCF and will likely continue to be so in the highly effective modulator era.

## Materials and methods

### Source of key chemicals and bacterial strains

|  | Source of materials | Catalog number |
| --- | --- | --- |
| **M-CSF** | Miltenyi | 130-096-491 |
| **Pan monocyte isolation kit** | Miltenyi | 130-096-537 |
| **3-oxo-C12-homoserinelactone (3OC12HSL)** | Sigma | O9139 |
| **C4-homoserinelactone (C4HSL)** | Chemodex | B0267 |
| **Ivacaftor** | Selleckchem | S1144 |
| **Tezacaftor** | Selleckchem | S7059 |
| **Elexacaftor** | Selleckchem | S8851 |
| **Seahorse mitostress kit** | Agilent |  |
| **THP-1 cells** | ATCC | TIB-202 |
| **PMA** | Sigma | P1585 |
| **PA14 WT** | (79) | Hogan Lab (DH122) |
| **PA14 $\Delta lasR$** | (80) | Hogan Lab (DH164) |
| **PA14 $\Delta lasR$ att::*PrhlI-lacZ*** | (81) | Hogan Lab |
| **DH1137 (LasR+)** | (82) | Hogan Lab (DH1137) |
| **DH1136 (LasR-)** | (82) | Hogan Lab (DH1136) |
| **DH2417 (LasR+), NC-AMT0101-1-2**<br>**(parent of NC-AMT0101-1)** | (82) | Hogan Lab (DH2417) |
| **DH2415 (LasR-), NC-AMT0101-1-1,**<br>**related to DH2417 with LasR**<br>**LOF (frameshift) allele** | (82) | Hogan Lab (DH2415) |

### THP-1 cell culture

THP-1 cells were cultured in RPMI with 10% FBS, 50 µM β-mercaptoethanol, and 50 μg/mL gentamicin in a 37°C humidity-controlled incubator with 5% CO_2_. The initial vial was expanded and frozen down in aliquots which were stored in liquid nitrogen with DMSO. Cultures from individual aliquots were maintained in vitro for ∼2 months prior to being discarded. THP-1 monocytes were differentiated into macrophages by addition of 50 nM Phorbol 12-myristate 13-acetate (PMA) for 48 hours prior to experiments.

### Gentamicin protection phagocytosis assays with THP-1 derived macrophages

THP-1 cells were differentiated at 5 x 10^5^/mL in 24-well plates. They were then washed with PBS and medium was replaced with identical medium without antibiotics. Overnight LB cultures of bacteria of indicated strains were sub-cultured 1:10 into LB for 60 minutes, centrifuged at 8000 x g for 2 minutes, washed twice with antibiotic-free THP-1 medium, then resuspended and bacterial density was determined by spectrophotometry at 600 nm. OD600 of 1.0 was empirically determined to represent 10^9^ bacteria/mL media. After dilution, bacteria were added to THP-1 cells at a multiplicity of infection (MOI) of 10 bacteria per macrophage. Plates were incubated at 37°C for 20 minutes, then cells were washed once with PBS, then THP-1 medium with 10x gentamicin (500 μg/mL) was added for 15 minutes. Cells were then washed twice with PBS at 4°C followed by lysis with 250 µL 0.1% Triton X-100. Lysates were diluted 1:3 in PBS and then 10 µL was then plated overnight onto LB agar plates for colony forming unit (CFU) determination. Colonies were counted the next day and bacteria per 10^6^ macrophages was calculated. MOI was also determined empirically from the stock bacterial solutions for each assay as an internal control. Average of at least three technical replicates per assay was then log-transformed and results were combined from at least 5 assays performed on separate days (n for each experiment listed in figure legends).

### Phagocytosis competition assay

PA14 Δ*lasR* with a *lacZ* reporter gene integrated at a neutral site on the chromosome was used along with PA14 wild type (WT). Assay was identical to above except fixed percentages of PA14 WT and the Δ*lasR* strain were used at 100/0, 90/10, 70/30, 50/50, 30/70, 10/90, and 0/100 such that the total MOI of both strains together was constant at 10. Lysates were plated onto LB agar plates with 5-bromo-4-chloro-3-indolyl-β-D-galactoside (X-gal) for blue-white colony discrimination. 100/0 samples were entirely white and 0/100 samples were entirely blue, confirming the fidelity of the reporter. Total PA14 WT and Δ*lasR* colonies were counted for each well. Each experiment was performed in triplicate and the means of technical replicates for each assay were log-transformed and combined.

### Human subjects

Our blood draw protocol has been approved by the Dartmouth Health Institutional Review Board. Subjects were recruited from our CF clinic population if they were 18-65 years old, F508del homozygous or heterozygous, at clinical baseline without symptoms of exacerbation, and had normal hemoglobin levels. NonCF control subjects were recruited by IRB-approved flyers; smokers or those with respiratory disease, immunologic disorders, or pulmonary medication use were excluded. Phlebotomy was performed (100 mL whole blood isolated) for monocyte isolation by our research nurse and taken directly to the laboratory for downstream processing.

### Monocyte derived macrophage (MDM) generation and phagocytosis

MDMs were generated according to previously published protocols (37). Briefly, monocytes were isolated from peripheral blood by Ficoll gradient separation. They were quantified and plated directly onto plates as per intended assay to avoid the need for detachment and re-plating. MDMs were generated by treatment with 100 ng/mL M-CSF for seven days in RPMI/10% FCS with 50 μg/mL gentamicin. Medium was exchanged every 2-3 days. After seven days of differentiation, MDMs were cultured in RPMI/10% FCS + gentamicin, and CFTR modulators were added for 48 hours prior to the experiment. CFTR modulators were replaced with all media exchanges. Gentamicin protection assay was performed as above. Three to four technical replicates per subject were averaged and log-transformed. Data on paired *lasR* mutants were combined with previously published data on *lasR*-WT strains (37).

### Seahorse extracellular flux assay

Monocytes were plated initially onto a Seahorse 96-well plate at 50,000 per well and differentiated into MDM directly in the wells for seven days as above. At that point, differentiation medium was removed and CFTR modulators were added for 48 hours. Pa-conditioned medium was prepared in advance by culturing bacteria overnight in Seahorse mitostress assay medium, normalizing cultures to an OD600 of 0.500, centrifuging to pellet bacteria then passing supernatant through a 0.22 µm filter. This was defined as 100% conditioned medium. Aliquots were then frozen at -20° C until use.

On the day of assay, cell culture medium was exchanged for mitostress assay medium as per manufacturer’s instructions. A mitostress assay with acute injection was run on an Agilent Seahorse XFe96 platform, with 20 µL conditioned medium in injection port A (final concentration 10%), plain mitostress medium was included for control wells. Oligomycin, FCCP, and Rotenone/Antimycin A were placed into ports B-D respectively. Four to eight technical replicates per condition were run. Parameters including response to injection, maximal respiration, and others were calculated as per manufacturer’s instructions. Data were combined with previously published results with a *lasR*-WT strain only (37).

### THP-1 Seahorse extracellular flux assay

THP-1 macrophages were differentiated directly on a Seahorse 96-well plate with 50 nM PMA at 50,000 cells/well for 48 h. Seahorse extracellular flux assay was then run with varying concentrations of 3OC12HSL or C4HSL in injection port A. Five to six technical replicates were run per condition for each assay and averaged. Five independent experiments were run.

### Cytokine analysis

MDMs were differentiated for seven days as above then treated for 48 hours with CFTR modulators or DMSO. Bacteria were prepared in an identical manner to the phagocytosis assays. After exchange for antibiotic-free media, MDMs were infected with indicated strains at an MOI of 10 for 2 hours, then washed and placed in media with gentamicin for an additional 2 hours. Supernatants were then collected, centrifuged at 14,000 x g at 4°C for 10 minutes, transferred to a new tube, and frozen at -20°C until needed. Samples were batched and cytokine multiplex analysis was run by the Dartmouth Immunology and Flow Cytometry Core. One replicate per condition was run. Samples with levels below the limit of the detection of the assay were set to a concentration of 0, and data were log-transformed prior to analysis.

### Figure generation and statistical analysis

Figures were generated with GraphPad Prism 10. Statistical analyses were performed separately in R, with mixed model linear analysis using nlme package version 3.1. Models include all variables of interest as fixed variables and assay date (for THP-1 experiments) or subject (for MDM experiments) as random variables.

## Acknowledgments

Funding for DSA provided by the CF Foundation (ARIDGI19B0, ARIDGI20L0, STANTO19R0). THH is supported by DartCF (NIH grant P30-DK117469) and Cystic Fibrosis Foundation Research Development Program (BOMBER24G0). AA is supported by the NIH (R56HL158923, R01HL174700) and CFF (ASHARE21G0). Dartmouth Immunology and Flow Cytometry Core is supported by NIH 5P30CA023108. The authors declare no competing interests and the funders had no direct role in the design or conduct of these studies.

