## Supplemental Figures 1-3 for "*Pseudomonas aeruginosa lasR* Mutants Resist Phagocytosis and Alter Inflammatory Cytokine Production by Cystic Fibrosis Macrophages"

### Slide 1
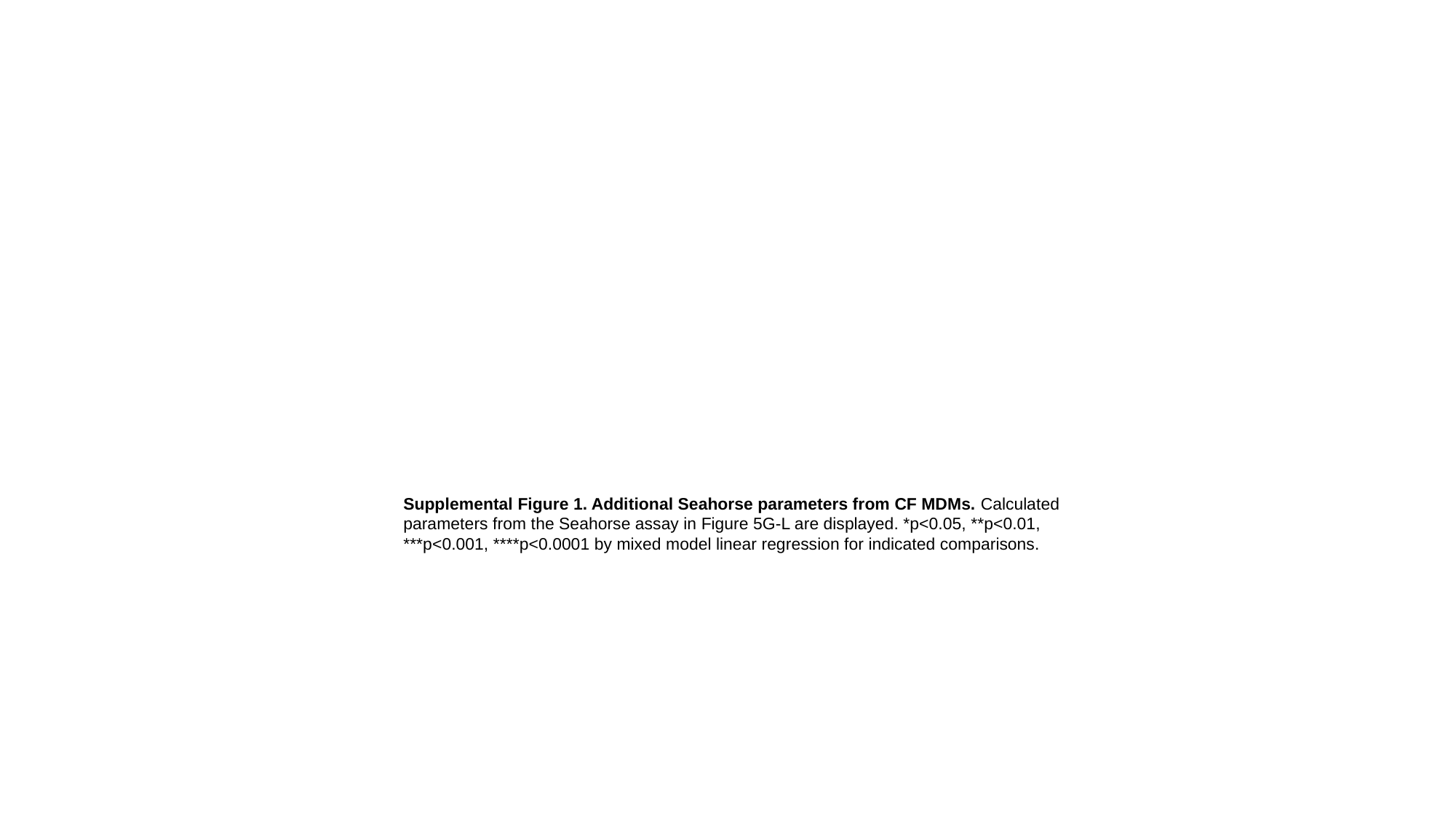

Supplemental Figure 1. Additional Seahorse parameters from CF MDMs. Calculated parameters from the Seahorse assay in Figure 5G-L are displayed. *p<0.05, **p<0.01, ***p<0.001, ****p<0.0001 by mixed model linear regression for indicated comparisons.

### Slide 2
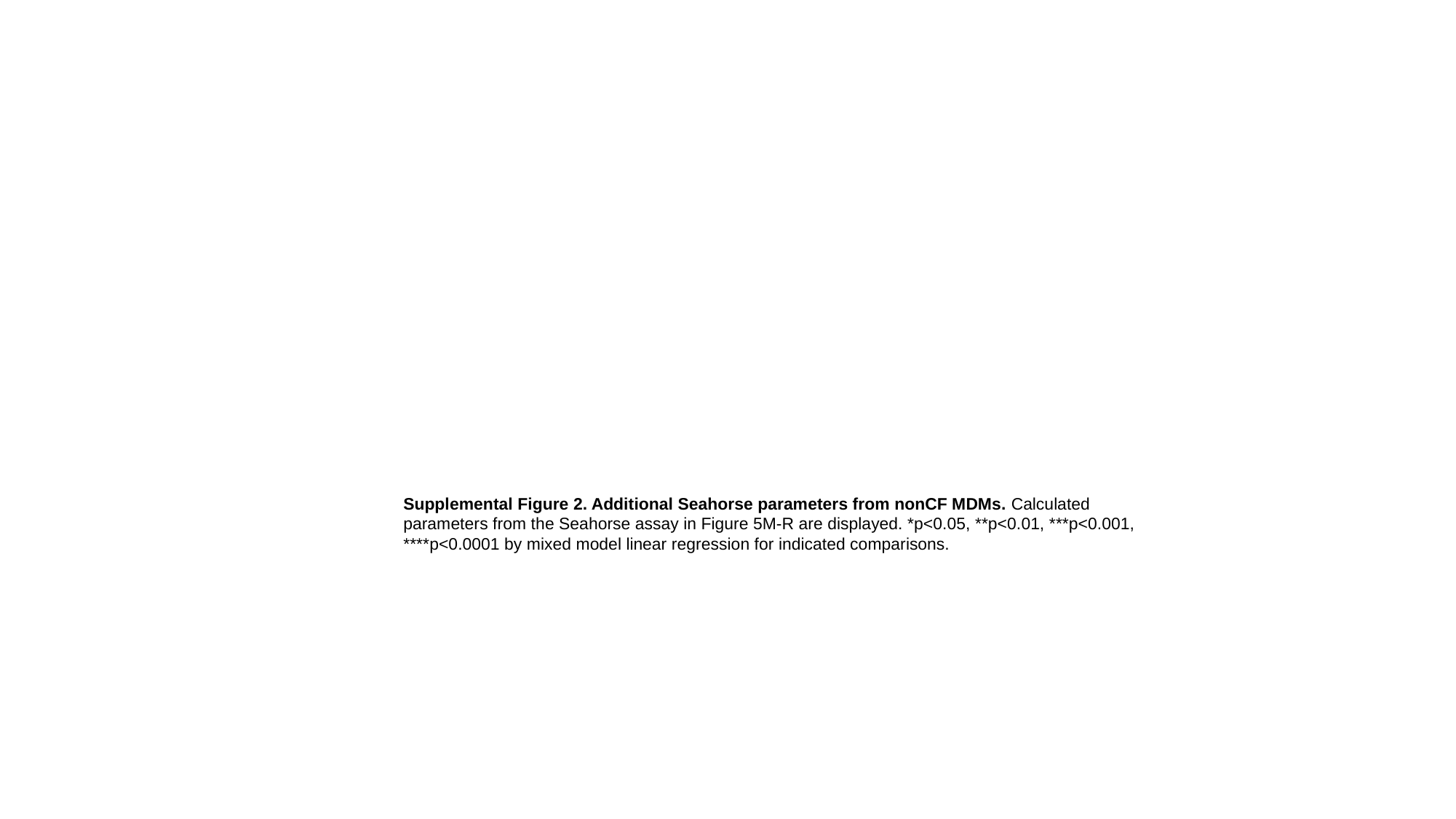

Supplemental Figure 2. Additional Seahorse parameters from nonCF MDMs. Calculated parameters from the Seahorse assay in Figure 5M-R are displayed. *p<0.05, **p<0.01, ***p<0.001, ****p<0.0001 by mixed model linear regression for indicated comparisons.

### Slide 3
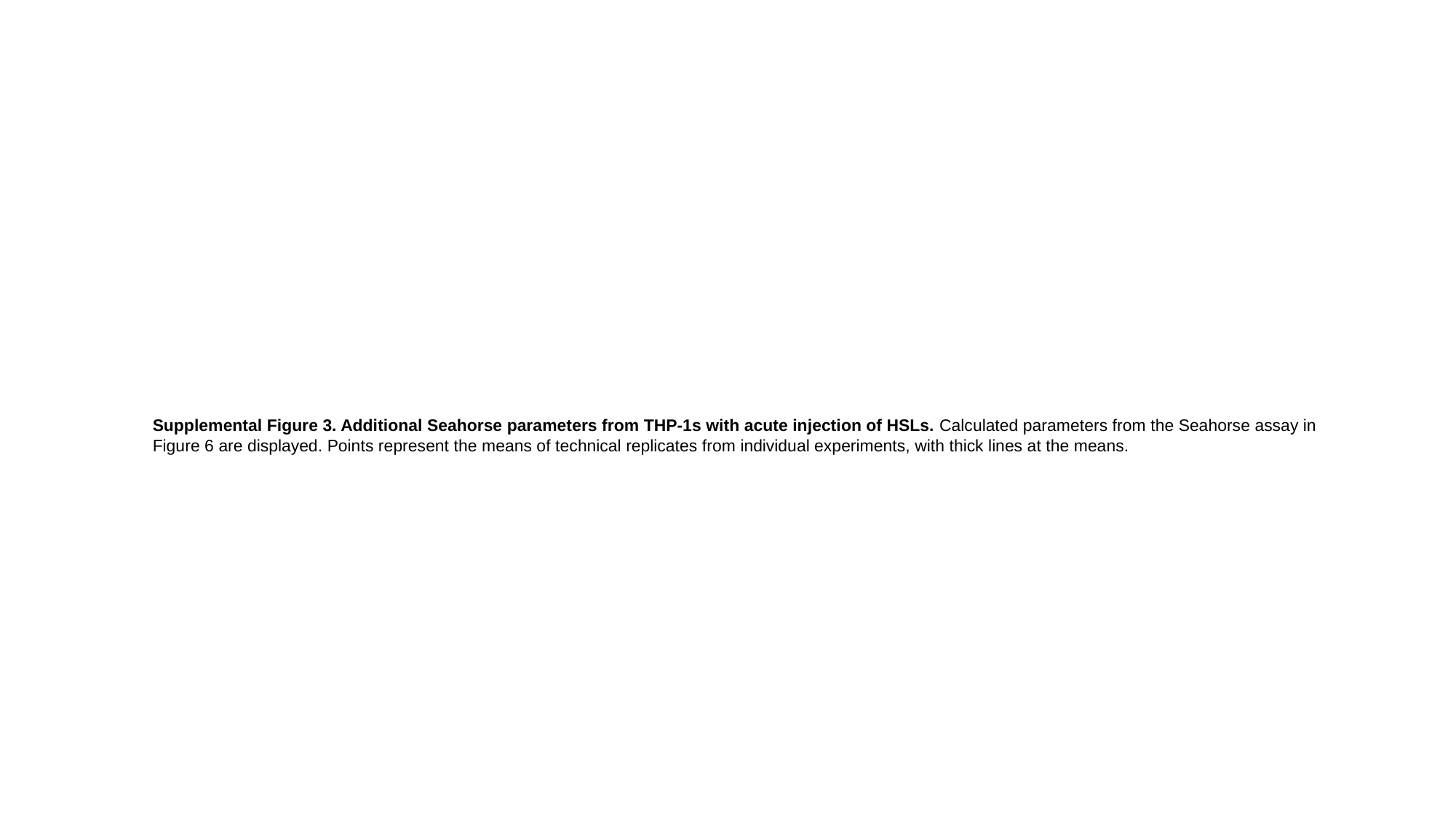

Supplemental Figure 3. Additional Seahorse parameters from THP-1s with acute injection of HSLs. Calculated parameters from the Seahorse assay in Figure 6 are displayed. Points represent the means of technical replicates from individual experiments, with thick lines at the means.
